# Resting-state network dysfunction in Alzheimer’s disease: a systematic review and meta-analysis

**DOI:** 10.1101/108282

**Authors:** AmanPreet Badhwar, Angela Tam, Christian Dansereau, Pierre Orban, Felix Hoffstaedter, Pierre Bellec

## Abstract

**INTRODUCTION:** We performed a systematic review and meta-analysis of the Alzheimer’s disease (AD) literature to examine consistency of functional connectivity alterations in AD dementia and mild cognitive impairment (MCI), using resting-state functional magnetic resonance imaging (rsfMRI).

**METHODS:** Studies were screened using a standardized procedure. Multiresolution statistics were performed to assess the spatial consistency of findings across studies.

**RESULTS:** Thirty-four studies were included (1,363 participants, average 40 per study). Consistent alterations in connectivity were found in the default-mode, salience and limbic networks in patients with AD dementia, MCI, or in both groups. We also identified a bias in the literature towards specific examination of the default-mode network.

**DISCUSSION:** Convergent evidence across the literature supports the use of resting-state connectivity as a biomarker of AD. The locations of consistent alterations suggest that metabolically expensive hub regions in the brain might be an early target of AD.

## 1. BACKGROUND

Alzheimer’s disease (AD), the leading cause of dementia in the elderly [1], exists on a continuum comprised of an asymptomatic preclinical stage, a middle stage of mild cognitive impairment (MCI), and a final stage of classic AD dementia. Symptoms usually start after the age of 65, except in the fewer than 5% of patients with early-onset autosomal dominant AD (ADAD) [2]. Drugs currently available for the treatment of AD provide only limited, short-term treatment of symptoms, without influencing the underlying pathological disease mechanism [3]. Trials of disease-modifying therapies for patients with AD dementia have been unsuccessful, likely because intervention at this stage is too late to affect the neurodegenerative process. The focus now is on therapeutic intervention at the pre-symptomatic stage (i.e. secondary prevention), with delay of dementia onset constituting a major clinical endpoint for clinical trials. This approach depends on the identification of biomarkers that can aid early diagnosis or characterization of this multifaceted disease, before its clinical expression [4]. Intrinsic brain connectivity, measured using resting-state functional magnetic resonance imaging (rsfMRI), is gaining popularity as an AD biomarker [4–6]. RsfMRI measures the intrinsic low-frequency fluctuations in the blood oxygen level-dependent (BOLD) signal, a proxy measure for neural processing in the brain, that can be used to identify spatially distributed functional connectivity networks [7]. Functional connectivity has been linked to the machinery of synaptic communication [8,9], which shows dysfunction early in the course of AD [5,6].

Altered connectivity in AD was initially reported in the default mode network (DMN) [10], spurring additional investigations in this network and, to a lesser extent, in other intrinsic connectivity networks (ICNs). To date, multiple studies have reported ICN disturbances in patients with AD dementia and MCI, pre-symptomatic ADAD mutation-carriers, and cognitively normal individuals carrying the at-risk APOEε4 allele and/or showing in-vivo evidence of amyloid deposition [11,12]. Despite such promising findings in support of the use of rsfMRI as an early biomarker of AD, the overall effect of AD on ICNs remains poorly characterized due to several limitations in the literature. The most prominent factor is that only a handful of studies systematically examine and report connectivity differences throughout the whole brain, while most studies focus on a single ICN. In addition, much of the published literature employ small sample sizes, and individually suffer from low statistical power. Finally, there may be systematic differences amongst studies that are difficult to assess, such as acquisition details, methods used to process and extract connectivity measures from fMRI, or the criteria applied for inclusion in each clinical group [13]. Our aim was to perform a systematic review and multi-resolution meta-analysis to examine the consistency of intrinsic connectivity alterations in MCI and late-onset AD (LOAD) dementia across the literature. We concentrated on the MCI and LOAD populations because the number of published studies was amenable to systematic meta-analysis.

## 2. METHODS

### 2.1 Literature Search

We conducted a systematic review of PubMed published studies up to December 3rd, 2015 in accordance with the PRISMA (Preferred Reporting Items for Systematic Reviews and Meta-Analyses) guidelines [14]. Search terms and combinations used are provided in Supplemental Table 1. Results were filtered for duplicates within each of the two main search categories (patients with AD dementia or MCI). Unique search results underwent further screening as described below.

**Table 1:** Characteristics of rsfMRI studies included in the meta-analysis. Studies investigating both Alzheimer’s disease (AD) dementia and mild cognitive impairment (MCI) cohorts are indicated by *, and those reporting significant coordinates for both AD and MCI patients, relative to matched healthy controls (HC), are indicated by **. In “**bold**” we indicate 7 studies using shared cohorts. Coordinates from these 7 studies were subsequently pooled under 4 studies (indicated by ^**a**^, ^**b**^, and ^**c**^), under the corresponding earliest publication using the cohort. In column “**Method**”, when both seed-based (SB) and independent component analysis (ICA) rsfMRI methods were employed by a study, the <u>underlined</u> method indicates the method associated with reported coordinates. For column “**Seed Region/ICN Investigated**”, all seed regions and ICN (intrinsic connectivity networks) investigated are listed, irrespective of significant findings. Abbreviations: a (anterior); B (Brucker); d (dorsal); f (female); GE (General Electrics)l (left); lat (lateral); m (male); med (medial); n (number of subjects); p (posterior); P (Philips); r (right); S (Siemens); sd (standard deviation); T (Tesla); v (ventral); ACC (anterior cingulate cortex); AG (angular gyrus); dlPFC (dorsolateral prefrontal cortex); ICC (isthmus of cingulate cortex); mPFC (medial prefrontal cortex); medialFC (medial frontal cortex); middleFC (middle frontal cortex); MrD (marginal division); PCC (posterior cingulate cortex); saVC (secondary associative visual cortex); AN (auditory network); ATN (attentional network); DMN (default mode network); ECN (executive control network); SMN (sensorimotor network); prVIS (primary visual network); sVIS (secondary visual network)

| Study | N | AD |  |  |  |  | HC |  |  |  |  | AD |  | Scanner | Method | Seed Region/ICN Investigated |
| --- | --- | --- | --- | --- | --- | --- | --- | --- | --- | --- | --- | --- | --- | --- | --- | --- |
|  |  | n | m | f | age | sd | n | m | f | age | sd | <HC | >HC |  |  |  |
| Wang et al. (2007) | 28 | 14 | 7 | 7 | 70.2 | 6.3 | 14 | 7 | 7 | 69.6 | 5.5 | x |  | 1.5 T S | SB | PCC |
| Zhang et al. (2009)<sup>a</sup> | 32 | 16 | 6 | 10 | 71.6 | 5.1 | 16 | 7 | 9 | 71.3 | 4.9 | x | x | 1.5 T P | SB | PCC |
| Zhang et al. (2010)<sup>a</sup> | 55 | 39 | 18 | 21 | 73.4 |  | 16 | 7 | 9 | 71.3 | 4.9 | x | x | 1.5 T P | SB | PCC |
| Sheline et al. (2010) | 83 | 35 |  |  |  |  | 48 |  |  |  |  | x | x | 3.0 T S | SB | precuneus |
| Zhou et al. (2010) | 24 | 12 | 5 | 7 | 63.3 | 7.7 | 12 | 5 | 7 | 62.0 |  | x | x | 1.5/3.0/4.0 T S/GE/B | SB & ICA | AG (l), pregenual ACC (r) |
| **Gili et al. (2011) | 21 | 11 | 7 | 4 | 71.9 | 7.9 | 10 | 7 | 3 | 64.1 | 10.5 | x |  | 3.0 T S | SB & ICA | PCC, mPFC |
| Wu et al. (2011)<sup>b</sup> | 31 | 15 | 6 | 9 | 64.0 | 8.3 | 16 | 7 | 9 | 65.0 | 9.2 | x |  | 3.0 T S | ICA | DMN |
| Li et al. (2012)<sup>b</sup> | 31 | 15 | 6 | 9 | 64.0 | 8.3 | 16 | 7 | 9 | 65.0 | 9.2 | x |  | 3.0 T S | ICA | ATN (d,v) |
| Damoiseaux et al. (2012) | 39 | 21 | 9 | 12 | 64.2 | 8.7 | 18 | 12 | 6 | 62.7 | 10.3 | x | x | 3.0 T GE | ICA | DMN (a,p,v), SMN |
| *Binnewijzend et al. (2012) | 82 | 39 | 23 | 16 | 67.0 | 8.0 | 43 | 23 | 20 | 69.0 | 7.0 | x |  | 1.5 T S | ICA | DMN, working memory (l,r), visuospatial attention (d), spatial attention (v), SMN, auditory-language, prVIS, sVIS, basal ganglia-cerebellum |
| Kenny et al. (2012) | 32 | 16 |  |  | 77.3 | 8.9 | 16 |  |  | 76.3 | 8.3 |  | x | 3.0 T P | SB | hippocampus (l,r), PCC, precuneus, prVIS |
| *Zhu et al. (2013) | 22 | 10 | 7 | 3 | 72.9 | 7.9 | 12 | 5 | 7 | 73.8 | 6.5 | x |  | 3.0 T GE | SB | ICC (l,r) |
| Balthazar et al. (2014) | 37 | 20 |  |  | 73.9 | 8.2 | 17 |  |  | 72.3 | 6.4 | x | x | 3.0 T P | ICA | DMN (d,v), SN (a,p) |
| *Yao et al. (2013)<sup>c</sup> | 62 | 35 | 12 | 23 | 72.4 | 8.5 | 27 | 16 | 11 | 69.2 | 6.5 | x |  | 3.0 T GE | SB | amygdala(l,r) |
| *Zhou et al. (2013)<sup>c</sup> | 62 | 35 | 12 | 23 | 72.4 | 8.5 | 27 | 16 | 11 | 69.2 | 6.5 | x | x | 3.0 T GE | SB | thalamus (l,r) |
| *Zhang et al. (2014)<sup>c</sup> | 62 | 35 | 12 | 23 | 72.4 | 8.5 | 27 | 16 | 11 | 69.2 | 6.5 | x |  | 3.0 T GE | SB | MrD (l,r) |
| Gour et al. (2014) | 28 | 14 | 6 | 8 | 75.1 | 2.9 | 14 | 4 | 10 | 72.8 | 3.0 | x | x | 3.0 T S | SB | PCC, perirhinal cortex (l,r), dlPFC (l,r) |
| Weiler et al. (2014) | 48 | 22 | 6 | 16 | 73.4 | 5.7 | 26 | 6 | 20 | 70.0 | 6.6 | x | x | 3.0 T P | SB | PCC, Wernicke's (l); Broca's (l), dlPFC (l,r), saVC |
| Balachandar et al. (2015) | 30 | 15 | 9 | 6 | 67.3 | 6.6 | 15 | 9 | 6 | 64.4 | 8.9 | x | x | 3.0 T S | ICA | DMN, thalamic, ECN |
| **Pasquini et al. (2015) | 43 | 21 | 8 | 13 | 72.3 | 8.6 | 22 | 6 | 16 | 66.3 | 9.0 | x | x | 3.0 T P | ICA | DMN (a,p) |
| Adriaanse et al. (2014) | 59 | 28 | 17 | 11 | 72.0 | 4.9 | 31 | 17 | 14 | 72.0 | 4.3 | x |  | 1.5 T S | ICA | DMN, VIS (med, lat), AN, SMN, ECN, dorsovisual (l,r) |
| **Yi et al. (2015) | 23 | 11 | 1 | 10 | 64.2 | 2.4 | 12 | 3 | 9 | 71.8 | 1.2 | x |  | 3.0 T GE | ICA | DMN, SN |
| Study | N | MCI |  |  |  |  | HC |  |  |  |  | MCI |  | Scanner | Method | Seed Region/ICN Investigated |
|  |  | n | m | f | age | sd | n | m | f | age | sd | <HC | >HC |  |  |  |
| Sorg et al. (2007) | 40 | 24 | 13 | 11 | 69.3 | 8.1 | 16 | 10 | 6 | 68.1 | 3.8 | x |  | 1.5 T S | ICA | VIS, AN, ATN (v), spatial attention, DMN |
| Bai et al. (2009) | 56 | 30 | 15 | 15 | 72.5 | 4.4 | 26 | 12 | 14 | 71.6 | 5.3 | x | x | 1.5 T GE | SB | PCC |
| **Gili et al. (2011) | 20 | 10 | 6 | 4 | 71.2 | 4.1 | 10 | 7 | 3 | 64.1 | 10.5 | x |  | 3.0 T S | SB & ICA | PCC, mPFC |
| Bai et al. (2011) | 44 | 26 | 19 | 7 | 71.4 | 4.3 | 18 | 10 | 8 | 70.3 | 4.7 |  | x | 1.5 T GE | ICA | DMN |
| Xie et al. (2014) | 56 | 30 | 19 | 11 | 72.6 | 4.8 | 26 | 14 | 12 | 70.3 | 4.8 | x |  | 1.5 T GE | SB | postcentral gyrus (l), hippocampus (l), medialFC (l), middleFC (l), precuneus (l,r), insula (l,r) |
| Jin et al. (2012) | 16 | 8 | 5 | 3 | 60.6 | 3.2 | 8 | 4 | 4 | 60.9 | 8.3 | x | x | 3.0 T GE | ICA | DMN |
| Han et al. (2012) | 80 | 40 | 7 | 33 | 86.3 | 4.5 | 40 | 15 | 25 | 86.3 | 4.5 | x | x | 1.5 T GE | SB | PCC |
| Liang et al. (2012) | 32 | 16 | 6 | 10 | 68.5 | 7.8 | 16 | 6 | 10 | 67.2 | 8.4 | x | x | 3.0 T S | SB | AG (l,r), supramarginal gyrus (l,r), intraparietal sulcus (r) |
| *Hahn et al. (2013) | 54 | 28 | 14 | 14 | 69.5 | 7.1 | 26 | 10 | 16 | 65.5 | 7.8 | x |  | 3.0 T P | ICA | DMN (a,p), ATN (d,v), ECN (l,r), SMN, VIS |
| Myers et al. (2014) | 35 | 23 | 14 | 9 | 69.3 | 7.4 | 12 | 5 | 9 | 63.8 | 5.2 | x |  | 3.0 T P | ICA | DMN (a,p), ATN (l,r, d), SN, prAN |
| Koch et al. (2015) | 40 | 24 | 14 | 10 | 68.2 | 8.4 | 16 | 7 | 9 | 64.8 | 5.4 | x |  | 3.0 T P | ICA | DMN (a,p), ATN (l,r,d), SN, prAN |
| **Pasquini et al. (2015) | 44 | 22 | 11 | 11 | 65.3 | 8.7 | 22 | 6 | 16 | 66.3 | 9.0 |  | x | 3.0 T P | ICA | DMN (a,p) |
| Das et al. (2015) | 69 | 30 | 14 | 16 | 71.6 | 6.8 | 39 | 18 | 21 | 70.6 | 9.0 | x |  | 3.0 T S | SB | hippocampal subregions |
| Gardini et al. (2015) | 42 | 21 | 13 | 8 | 70.6 | 4.7 | 21 | 7 | 14 | 69.8 | 6.5 |  | x | 3.0 T GE | SB | PCC, mPFC |
| **Yi et al. (2015) | 32 | 20 | 4 | 16 | 71.0 |  | 12 | 3 | 9 | 71.8 | 1.2 | x | x | 3.0 T GE | ICA | DMN, SN |

### 2.2 Study Selection

#### 2.2.1 Inclusion/Exclusion Criteria

Search results were subjected to two successive rounds of screening with increasingly stringent criteria. The initial screen was performed on the abstracts of search results. An article was included if the abstract indicated it was a peer-reviewed, original research article written in English, and employed rsfMRI to study LOAD and/or MCI in humans. Review articles, case reports, letters to editors, and studies with subjects in whom MCI was associated with other diseases were omitted.

Following the initial screening, we applied the following inclusion criteria: (a) used seed-based or Independent Component Analysis (ICA) rsfMRI methods, (b) investigated functional connectivity between patients (AD dementia or MCI) and age-matched healthy controls (HC), and (c) reported peak coordinates of significant statistical differences in average connectivity between groups, as well as the direction of difference.

### 2.2 Data Extraction

One reviewer (AB) conducted the search using the various search terms and screened for duplicates. Two reviewers (authors AB and AT) independently screened all unique search results for potential inclusion in the meta-analysis. Only articles passing both reviewers’ approval were considered for final inclusion. For each ‘included’ article, coordinate data of significant between-group comparisons, such as AD versus HC, was transcribed by one of the two reviewers, and checked by two others (second reviewer and FH).

### 2.3 Meta-Analysis

We performed complementary network- and voxel-based quantitative meta-analyses on six main group comparisons: pooled AD dementia + MCI (ADMCI)<HC, ADMCI>HC, MCI<HC, MCI>HC, AD<HC, and AD>HC. While voxel-based meta-analysis has finer spatial resolution for findings with high anatomical consistency, we assumed the network-based approach would have better sensitivity for detecting consistent involvement of anatomically distributed networks. Coordinates from articles that used the same cohort were pooled under the PMID of the earliest publication and therefore treated as results from a single study to avoid counting a cohort more than once. It should be noted that for the remainder of our paper an individual article will be referred to as a “study”, and a group comparison or single analysis yielding network and/or localization information (e.g. ADMCI<HC) as a “contrast”.

#### 2.3.1 Network-based Statistics

We performed network-based statistics on seed coordinates (seed statistics) to assess whether seed regions were preferentially selected from within certain networks in the literature. We also performed network-based statistics on coordinate data of significant contrasts (contrast statistics) to assess the consistency of network-level findings in the AD literature. In particular, we performed three types of contrast statistics: (a) all coordinates irrespective of seed network; and, given the focus on the DMN in the literature, (b) coordinates associated with seeds inside the DMN only; and, (c) coordinates associated with seeds outside the DMN, i.e. non-DMN seeds. All analyses were conducted using a multi-resolution atlas of group-level functional brain parcellations derived from an independent rsfMRI dataset, the Bootstrap Analysis of Stable Clusters (BASC) - Cambridge atlas (https://dx.doi.org/10.6084/m9.figshare.1285615.v1). This atlas consists of nine functional parcellations capturing successively finer levels of spatial detail, of which we used parcellations at two resolutions: the first comprised of 7 commonly used, large-scale networks (R7 atlas), and the second containing 36 networks (R36 atlas). We used R7 and R36 atlases for contrast statistics, and only the R7 atlas for seed statistics. Since seeds were assigned indirectly for studies where coordinates were not provided, indirect assignment could not be performed with sufficient precision to use the R36 atlas. Assignment of seeds to one of the R7 networks was based on published coordinates, when available. When only anatomical labels were provided for seed regions, network assignment was based on (a) the centers of gravity in MNI space, when available, or (b) visual approximation if no further information was available. For ICA-based studies, network assignment was based on (a) network coordinates when provided, or (b) visual assignment to one or more of the seven networks based on the degree of spatial overlap.

We tested the spatial consistency of both seed and peak locations using the following approach. For each study we computed the number of coordinates falling within each network, after conversion of Talairach space coordinates into MNI space using the Lancaster transform [15], when necessary. Coordinates falling outside of the gray matter mask (ICBM152) were assigned to the closest network. To remain unbiased to the number of coordinates reported per study, we computed the ratio of coordinates falling within each network to the total number of coordinates reported per study. This ratio was then averaged across studies. Significance of findings was assessed using Monte Carlo permutation tests. Using the total number of coordinates per study, we generated a random assignment of coordinates to networks, taking into consideration the volume of each network. Coordinate counts per network were normalized as described above, followed by an averaging across studies. This Monte-Carlo sampling process was repeated 10,000 times. Thereafter, we compared the distribution of the average frequency obtained from the random sampling to the frequency obtained from the meta-analysis, resulting in p-value estimates [16]. Multiple comparisons across networks were accounted for using a false-discovery rate procedure (FDR q<0.05) [17]. The p-values below 0.05 that did not survive multiple comparisons were deemed as “trends”.

#### 2.3.2 Voxel-based Statistics

Voxel-level statistical analysis was performed using Activation Likelihood Estimation (ALE), a widely-used algorithm for coordinate-based meta-analysis of neuroimaging studies. ALE aims at delineating brain regions with above-chance convergence of reported coordinates across experiments [18]. Coordinates falling outside the gray matter mask were removed from analysis. We used the in-house ALE algorithm implementation in Matlab 8.3(.0.532), which treats each of the coordinates in a given experiment as a 3D Gaussian probability distribution centered at the given coordinate. The probability distributions acknowledge the spatial uncertainty associated with each experiment. For any given study, the width of the spatial uncertainty of its coordinates is determined based on empirical data on the between-subject and between-template variance representing the main components of this uncertainty [18]. Then, the probability distributions of all coordinates per included study are combined for each voxel, generating a modeled activation (MA) map. To limit the effect of multiple coordinates very close to one another within a given study, we used the ‘Non-Additive’ approach, which calculates MA maps by taking the maximum probability across overlapping Gaussians [18]. ALE scores were computed on a voxel-by-voxel basis by taking the union across these MA maps. To distinguish between ‘true’ and random convergence between studies (i.e., noise), ALE scores were compared to a null-distribution reflecting a random spatial association between experiments (10,000 permutations). Non-parametric p values were assessed at a familywise error (FWE) corrected threshold of p < 0.05 on cluster-level (cluster-forming threshold: p < 0.001 at voxel level) and transformed into t scores for display purposes. Only contrasts including more than 18 experiments were considered, as recommended in a recent large-scale simulation study [19].

## 3. RESULTS

### 3.1 Search Results

The results of the initial reference search and study exclusion for rsfMRI meta-analyses are presented in Figure 1. Thirty-four studies totaling 1,363 subjects (post-pooling of identical cohorts) met our inclusion criteria and were included in the meta-analysis (see Supplemental Table 2). The total included 352 MCI, 378 AD dementia (specifically LOAD) and 633 HC. The bulk (54%) of the studies had 20 or fewer subjects per group. Twenty studies (66.7%) investigated rsfMRI connectivity measures with other domains, cognition being most frequent, and some with levels of amyloid burden using PIB (Pittsburgh compound B), brain atrophy, and structural connectivity. Table 1 provides additional characteristics of the rsfMRI studies included in our meta-analysis, including scanner make, scanner strength, and seed region and/or ICN investigated.

**Figure 1:**
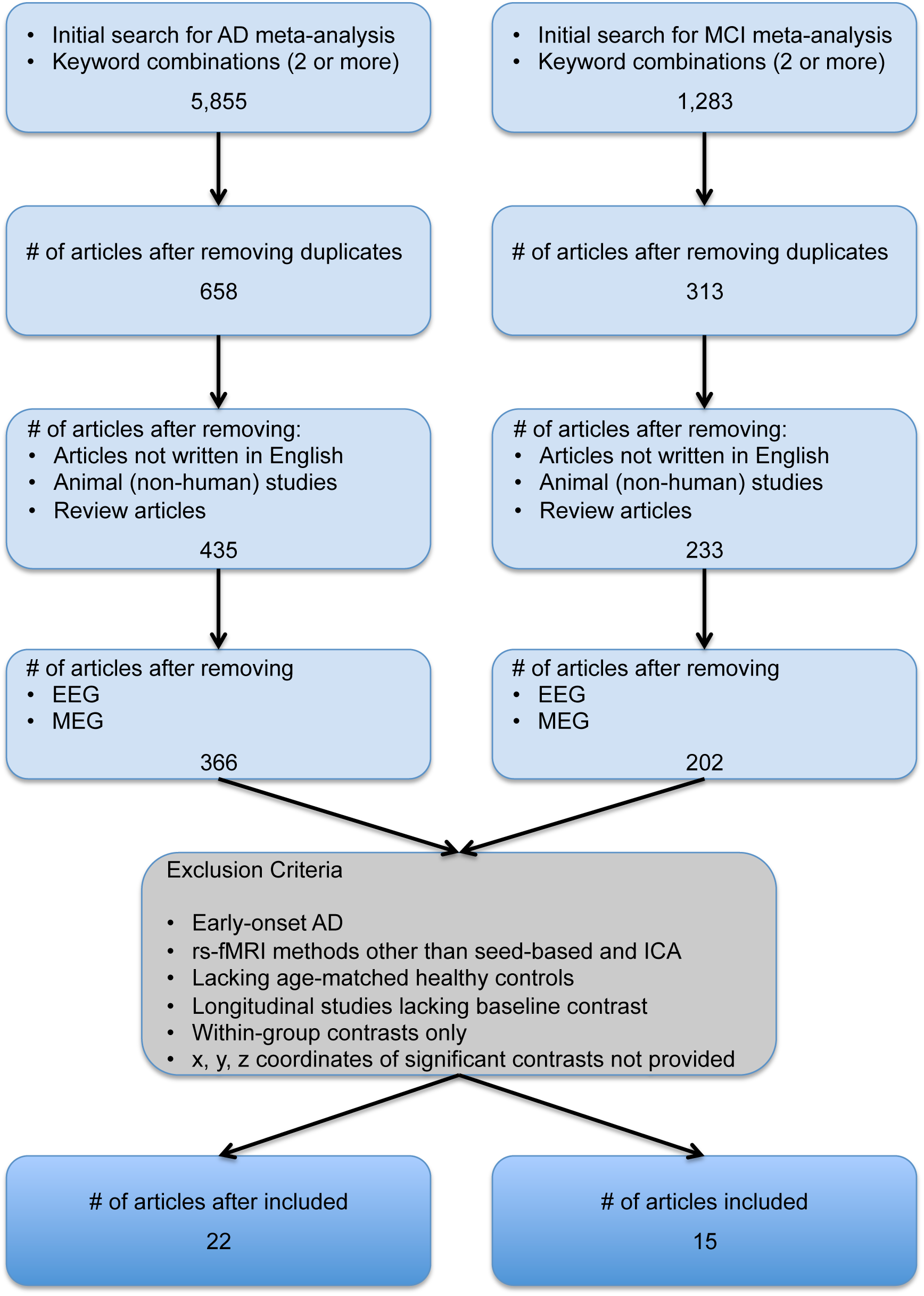
Flowchart of the study selection process. Selection process for AD and MCI studies included in the meta-analyses.

### 3.2 Network-Based Meta-analysis

#### 3.2.1 Seed Statistics

Using network-level statistics, we demonstrated a significant bias in the literature for ‘seed regions’ to originate within the DMN (Fig. 2), irrespective of the population (ADMCI, MCI or AD) being studied.

**Figure 2:**
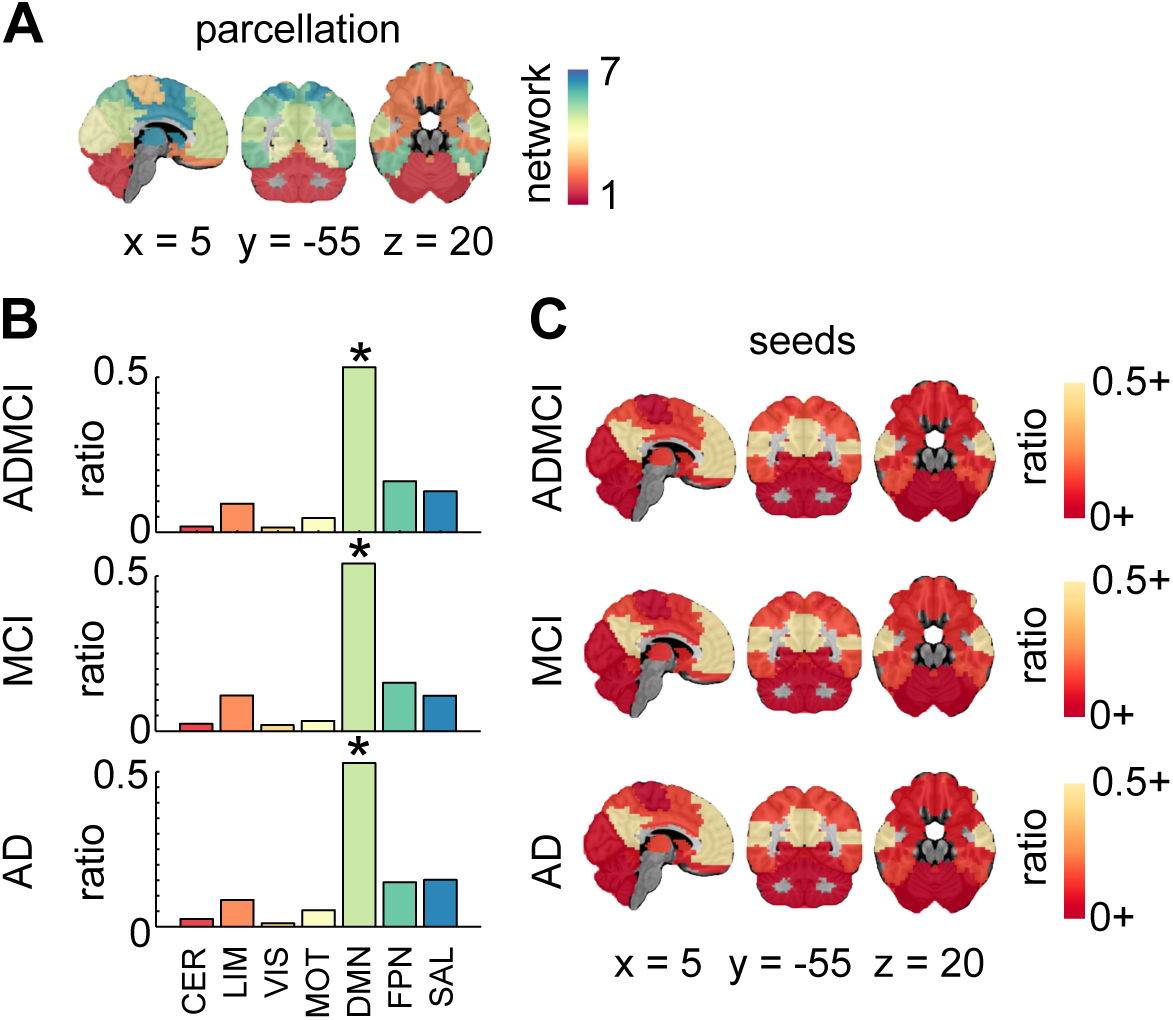
Seed region network-level findings. (A) R7 atlas; (B) Histograms show the ratio of counts (or hits) across the 7 networks for all seeds in ADMCI, MCI and AD. Significant (* denoting qFDR<0.05) prevalence of seeds in the DMN was demonstrated across all three cohorts; (C) Seed region hit maps (ratio of dysconnectivity coordinates in each network) at R7. Maps are superimposed onto the anatomical ICBM 152 template. x, y, and z MNI coordinates are given for sagittal, coronal and axial slices. Abbreviations: AD (Alzheimer’s disease dementia); MCI (mild cognitive impairment); HC (healthy control); CER (cerebellar network); LIM (limbic network); VIS (visual network); MOT (motor network); DMN (default-mode network); FPN (frontoparietal network); SAL (salience network)

#### 3.2.2 Contrast Statistics

We first examined R7 network-level statistics, and all seeds combined. Aberrant functional brain connectivity was observed in ADMCI, MCI, and AD, relative to HC (Figure 3). In the ADMCI cohort, we found both significant hypo- and hyper-connectivity in the DMN. Significant hyper-connectivity in the DMN and limbic (LIM) networks was observed in the MCI cohort. There was also a significant hypo-connectivity in the DMN for the AD group, and as a trend for the MCI group.

**Figure 3:**
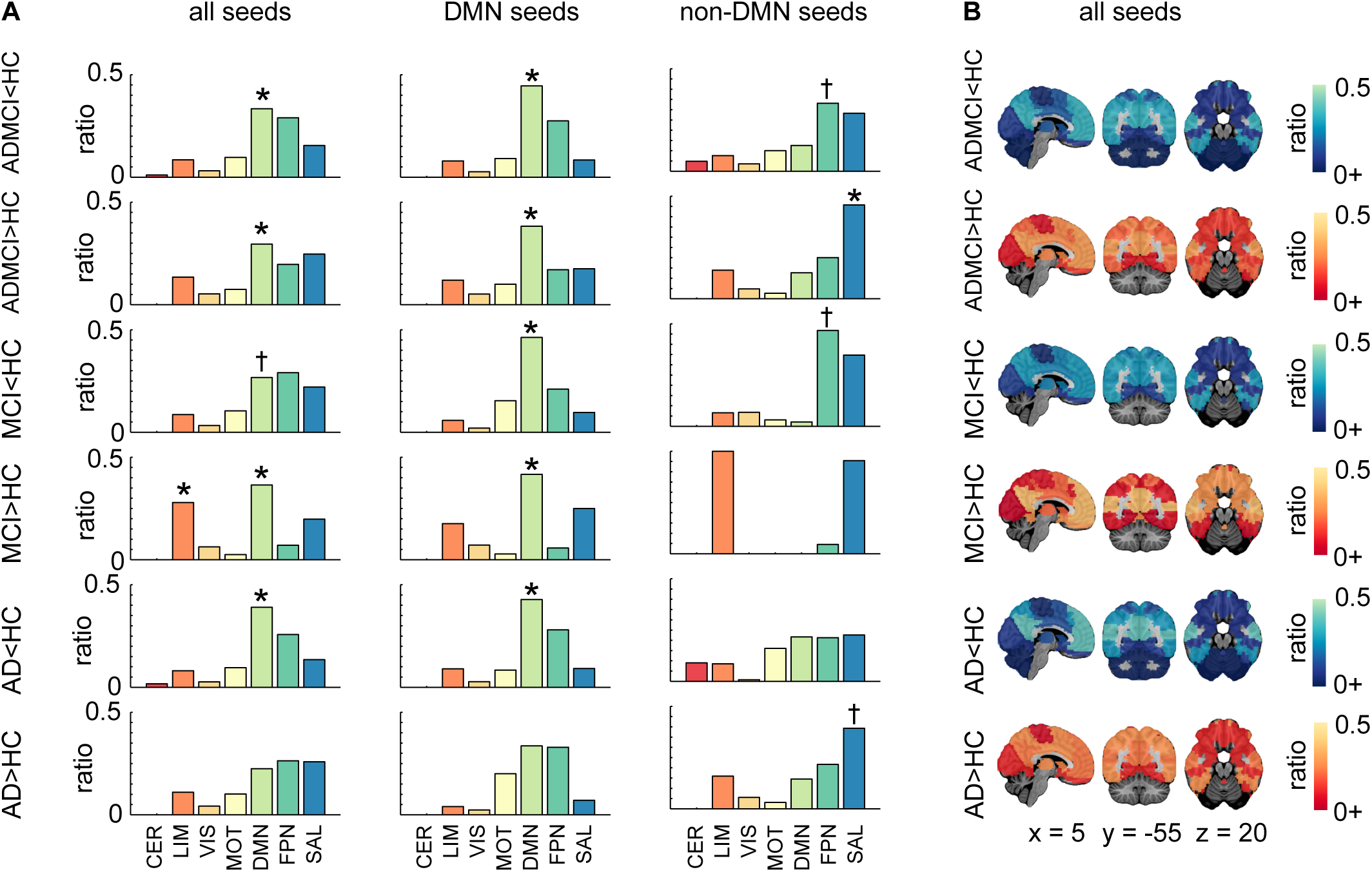
Network-level findings using the R7 atlas. (A) Histograms show per contrast the ratio of hits across the 7 networks for all seeds, DMN seeds only, and non-DMN seeds. Networks with significant count (or hit) ratios are indicated by * denoting qFDR<0.05, while † denotes p<0.05 uncorrected (B) Hit maps at R7 are shown for contrasts ADMCI<HC, ADMCI>HC, MCI<HC, MCI>HC, AD<HC, and AD>HC; Maps are superimposed onto the anatomical ICBM 152 template. x, y, and z MNI coordinates are given for sagittal, coronal and axial slices. Abbreviations: AD (Alzheimer’s disease dementia); MCI (mild cognitive impairment); HC (healthy control); CER (cerebellar network); LIM (limbic network); VIS (visual network); MOT (motor network); DMN (default-mode network); FPN (frontoparietal network); SAL (salience network)

We then refined the spatial localization of effects found in R7 using the R36 atlas. Significant DMN hypo-connectivity in AD and ADMCI cohorts was detected in the precuneus (PCu) and posterior cingulate cortex (PCC) (Figure 4). A trend for DMN hyper-connectivity was also observed in the PCu for ADMCI, and in both the PCu and PCC in MCI (Figure 4). The LIM hyper-connectivity was observed as a trend in the hippocampus and entorhinal cortex in MCI patients (Figure 4).

**Figure 4:**
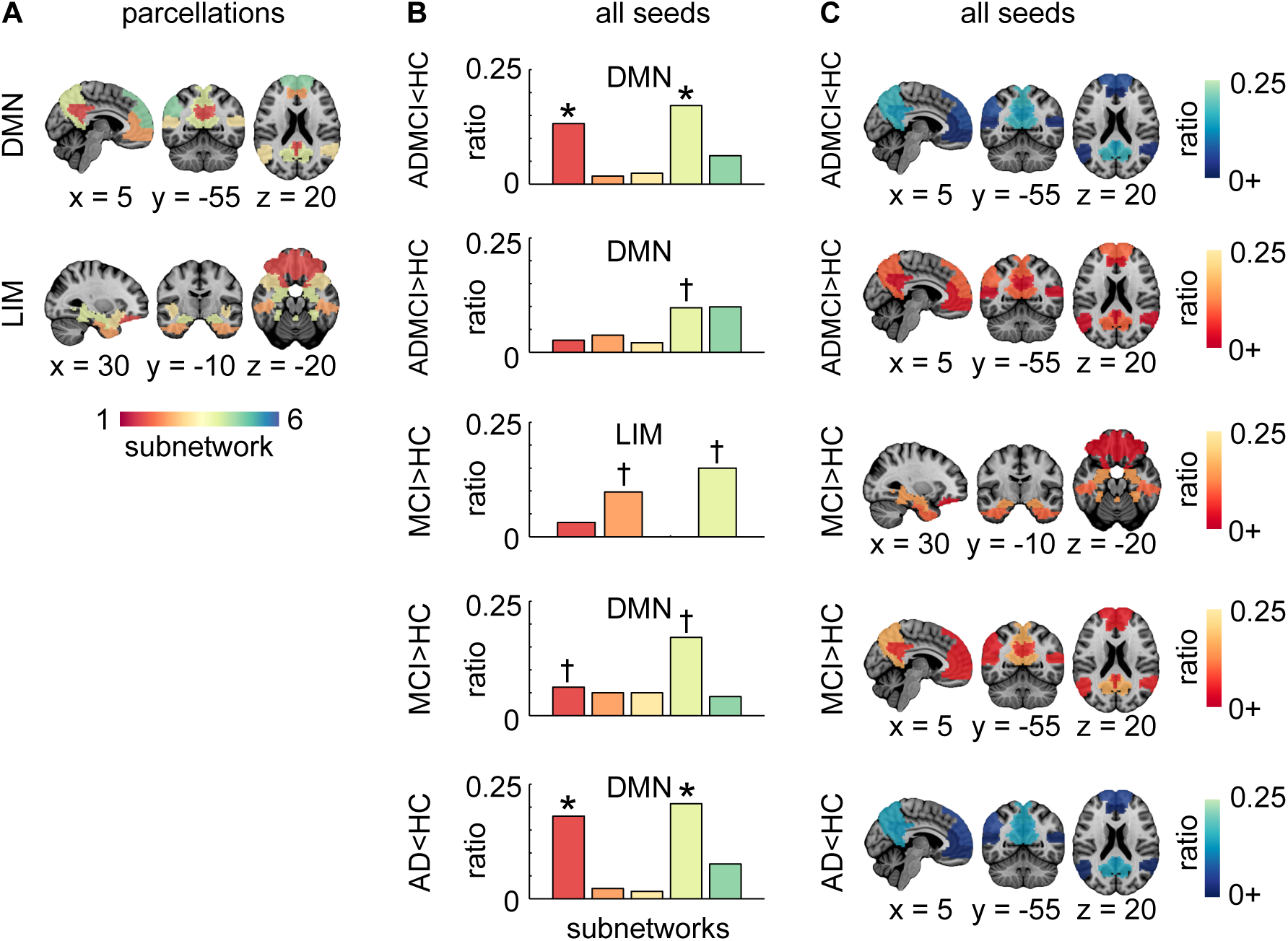
Network-level findings using the R36 atlas. (A) Functional template at R36 showing the breakdown of the DMN and LIM into subnetworks. These two networks were significant (qFDR<0.05 for contrasts ADMCI<HC for DMN, ADMCI>HC for DMN, MCI>HC for DMN and LIM, AD<HC for DMN) or trended towards significance (p<0.05 uncorrected for contrasts MCI<HC for DMN) for the ‘all seeds’ condition at R7; (B) Histograms show per selected contrast (as described in A), the ratio of counts (or hits) across the subnetworks. Subnetworks with significant hit ratios are indicated by * denoting qFDR<0.05, while † denotes a trend with p<0.05 uncorrected; (C) Hit maps at R36 for brain regions that overlap with significant or trending towards significance networks (as described in A). Maps are superimposed onto the anatomical ICBM 152 template. x, y, and z MNI coordinates are given for sagittal, coronal and axial slices. Abbreviations: AD (Alzheimer’s disease dementia); MCI (mild cognitive impairment); HC (healthy control); CER (cerebellar network); LIM (limbic network); VIS (visual network); MOT (motor network); DMN (default-mode network); FPN (frontoparietal network); SAL (salience network)

We finally investigated the robustness of findings with respect to the selection of seeds (DMN, non-DMN, or all combined), using the R7 atlas. Significant network-level findings derived from all seeds combined, as reported above, replicated when using DMN seeds alone (Figure 3A). In addition, a trend towards hypo-connectivity in MCI became significant using DMN seeds only. When focusing on non-DMN seeds studies, no significant effects were observed in the DMN, as expected. The only significant result was hyper-connectivity of the salience network (SAL) in ADMCI, also present as a trend in AD subjects.

### 3.3 Voxel-Based Meta-analysis

ALE results demonstrated significant hypo-connectivity in the PCC and PCu in the ADMCI and AD studies (Fig. 5, Supplemental Table 3), consistent with our network-level findings using R7 and R36 atlases. This observation was made both for all seeds combined, and DMN only seeds (Fig. 5, Supplemental Table 3).

**Figure 5:**
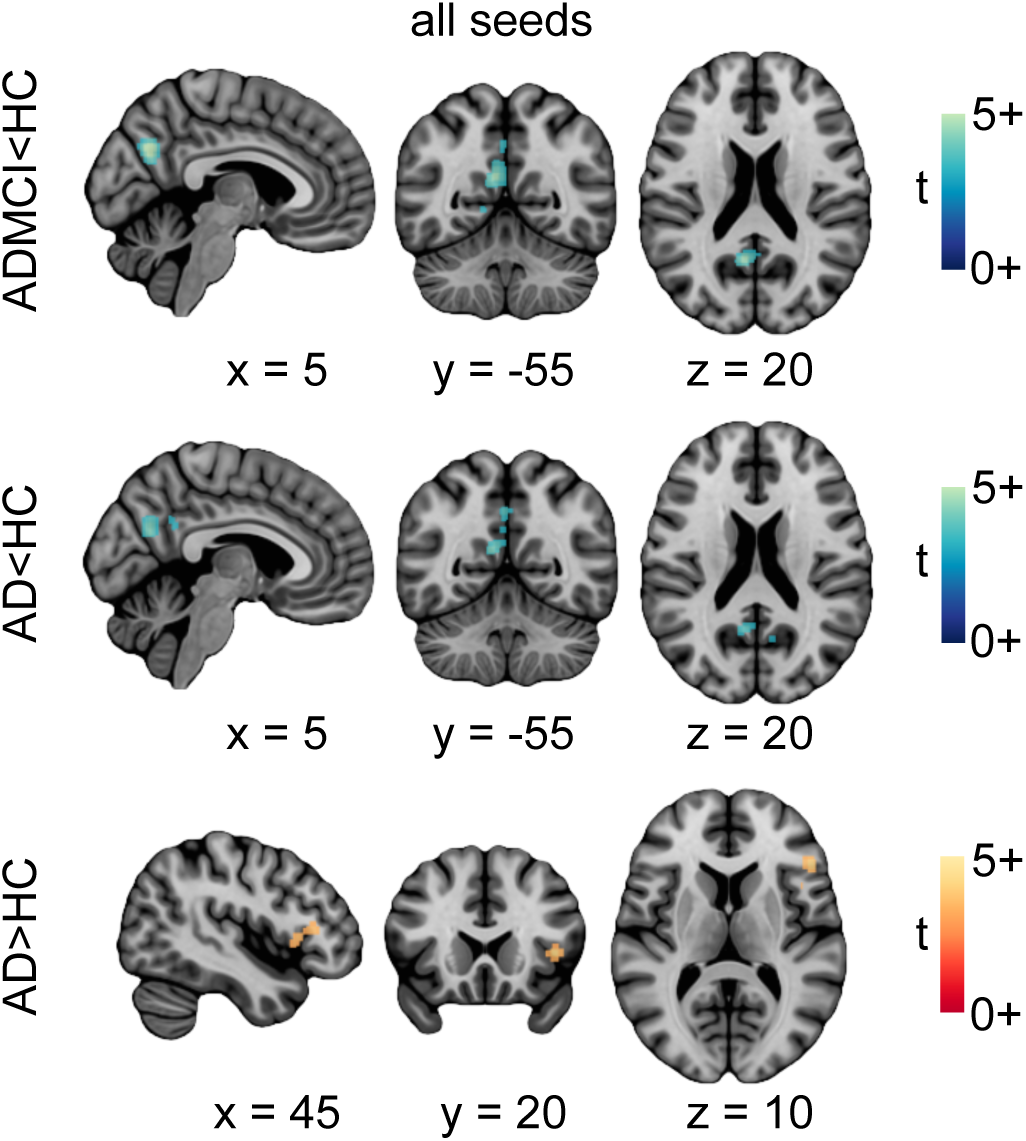
Location of significant convergence of the voxel-level findings. Regions’ exhibiting significant rsfMRI abnormalities for contrasts ADMCI<HC, AD<HC, and AD>HC. ALE images were thresholded at p < 0.05 (cluster-level FWE corrected for multiple comparisons; cluster-forming threshold p < 0.001 at the voxel-level) and displayed as t scores, with hyper-connectivity in red-orange and hypo-connectivity in blue-green. Maps are superimposed onto the anatomical ICBM 152 template. x, y, and z MNI coordinates are given for sagittal, coronal and axial slices. Abbreviations: AD (Alzheimer’s disease dementia); MCI (mild cognitive impairment); HC (healthy control)

Unlike the network-level analysis, using ALE we found diminished connectivity in the primary visual cortex, both in ADMCI and AD. This was observed for all seeds combined as well as DMN only seeds in ADMCI, and for DMN only seeds in AD. Finally, significant hyper-connectivity was observed in AD in the anterior insula (aINS) (Fig. 5, Supplemental Table 3), consistent with the trend in the LIM network observed with R36.

## 4. DISCUSSION

Our meta-analysis assessed the convergence of rsfMRI brain connectivity dysfunction findings in LOAD across a series of studies systematically selected from the literature. We implemented a multi-resolution approach to address the need for finer-grained localization of ICN abnormalities within the commonly-used larger networks [20]. We demonstrated that, despite a DMN bias in the literature, consistent intrinsic connectivity alterations in LOAD could be identified both within and outside of the DMN.

### 4.1 Connectivity changes in the Default Mode Network

We now discuss our findings in relation to prior relevant meta-analyses, as well as observations in early-onset AD, and in APOEε4 carriers. These cohorts were not included in our meta-analysis due to an insufficient number of available studies, but the literature is briefly reviewed.

#### 4.1.1. Late-onset AD

Our results revealed a consistent decrease in DMN connectivity in the ADMCI and AD cohorts, in particular the PCu and PCC. This finding is in line with two previous meta-analyses centered on the DMN [20,21]. The observation of DMN deterioration appears to be robust to the choice of analytical approaches, given that the former meta-analyses largely included studies using regional rsfMRI analysis methods, such as regional homogeneity and amplitude of low-frequency fluctuation.

Unlike our robust findings in AD subjects, DMN hypo-connectivity in MCI could only be demonstrated using network-level statistics on the subset of coordinates associated with DMN seeds. The trend for diminished connectivity localizing to the PCu when all coordinates were combined, together with a lack of voxel-level findings, suggests a weaker, more distributed network effect in MCI. Using a connectome-wide association approach, we recently reported pronounced widespread decreases in DMN connectivity in a large multisite MCI cohort [13]. The modest findings of our current literature-based meta-analysis may also be due to a lack of statistical power from having multiple, small, single-site samples. The clinically heterogeneous nature of this population might also have played a role, i.e. only a subset of MCI patients develop AD dementia [22], and there may be pathological subtypes as well for those who do [23]. We also demonstrated DMN hyper-connectivity in MCI and ADMCI using network-level statistics. Interpreting the directionality of changes in rsfMRI connectivity is difficult and may reflect compensation, as well as direct or indirect pathological mechanisms [24].

#### 4.1.2 Early-onset AD

DMN hypo-connectivity of similar magnitude to LOAD was demonstrated in early-onset non-autosomal-dominant AD [25,26], while in ADAD, DMN hypo-connectivity was slightly more pronounced than in LOAD [27]. Altered DMN connectivity was observed in asymptomatic adult mutation-carriers (PSEN1, PSEN2, or APP) with an “expected years to symptom onset” (eYO) of 12 years, on average [28,29], and a trend was detected in children with an eYO of up to three decades [30]. These findings suggest that altered functional connectivity may be a very early biomarker for AD.

#### 4.1.3 Cognitively normal individuals at genetic risk for LOAD

Altered DMN connectivity has been reported in cognitively normal APOEε4 carriers compared to non-carriers. These alterations were found across all age groups, i.e. elderly [12,31–33], middle-aged [34–36] and young adults [37,38]. In addition, connectivity changes in carriers were associated with lower cognitive performance in the middle-aged and elderly cohorts [32,34,36]. Interestingly, the connectivity changes in carriers occurred in the absence of PIB-detectable amyloid deposition [12,37,38].

### 4.2 Connectivity changes outside of the DMN

Despite the DMN bias in the literature, our meta-analysis confirmed that intrinsic connectivity disruptions in LOAD are not confined to the DMN. Across analyses, we found increased connectivity in the SAL in ADMCI and AD localized to the right aINS. Abnormal SAL connectivity was reported in ADAD [27] and APOEε4 carriers [33,34], though intensified connectivity was only detected in the latter. With a key hub in the aINS, the SAL plays a pivotal role in network switching between the DMN and frontoparietal network (FPN), two networks exhibiting competitive interactions during cognitive information processing [39]. Association of heightened SAL connectivity with reduced DMN connectivity in AD suggests that progressive DMN impairment is deleterious to SAL functioning [40].

We also found increased connectivity in the LIM in MCI. To date, heightened LIM connectivity has been reported in early-onset, non-autosomal dominant AD patients [26], and in individuals with subjective memory impairment [41]. The effect of APOEε4 carriage on LIM connectivity, however, lacks consensus [42–44]. Since LIM hyper-connectivity in early-onset AD patients was shown to correlate positively with memory performance, it is likely that increased connectivity in this network contributes to preserving function in the face of medial temporal lobe pathology [26].

### 4.3 Selective vulnerability of multimodal networks in AD

The DMN, SAL and FPN are multimodal networks that interconnect cortical regions associated with various cognitive functions, and they have been demonstrated computationally to support integrative information processing at the cost of being vulnerable to early and fast spreading of insults [45]. Supporting this theoretical finding is the recent observation that tau and amyloid-beta, despite their independent patterns of spatial deposition, show spatial overlap with brain tissue loss in hub regions of multimodal networks [46]. These multimodal networks are also metabolically expensive, and display higher rates of cerebral blood flow, aerobic glycolysis, and oxidative glucose metabolism [47]. The high value/high cost characteristics of the DMN, SAL and FPN make them vulnerable to (a) AD-associated pathogenic processes, such as metabolic dysfunction/oxidative stress, and (b) accumulation of toxic proteins, such as amyloid-beta [47]. Overall, our meta-analytic findings provide weight to the hypothesis that multimodal networks/regions maintaining reconfigurable connections to other brain areas are particularly susceptible to AD-associated neurodegeneration.

### 4.4 Limitations

While our literature search was systematic and exhaustive, we did not find an abundance of rsfMRI literature in AD and MCI cohorts, which clearly expresses the need for additional research. The relatively low number of experiments that met our inclusion criteria might have underpowered our voxel-level findings, especially for the MCI contrasts. In addition, our search demonstrated that typical studies featured low sample size, and also that analytical methods were quite variable in the field (a main reason for excluding a paper was due to methodology used). This setting is particularly amenable to questionable research practices, including “p-hacking” (testing several methods, reporting only one). Given the near absence of negative results reporting in the field, on one hand, and the large size of the rsfMRI field, on the other hand, there is no question that some amount of publication bias is also present. Meta-analytical tools such as funnel plots are available to detect both selective reporting and “p-hacking”, but are not feasible given current reporting practices in the rsfMRI community [48].

Another limitation of our study is the heterogeneity present in the included experiments, both in terms of population recruitment, scanning parameters, and processing choices [19,49]. There is also little doubt that the prominence of the DMN in our results is at least in part a reflection of the systematic bias of the literature towards that network, which we demonstrated quantitatively. The current trend towards large public samples is enabling unbiased meta-analyses, that pool neuroimaging data across many studies. This type of meta-analysis is becoming increasingly common in rsfMRI [13,50], and will hopefully resolve most of the aforementioned limitations in the future. Pooling of neuroimaging data will also facilitate proper investigations into the reported sex differences in AD (higher female prevalence).

### 4.5 Conclusions

Our meta-analysis demonstrated consistent connectivity alterations in the default-mode, salience and limbic networks in the spectrum of LOAD, supporting the use of resting-state connectivity as a biomarker of AD.

## ACKNOWLEDGEMENTS

The computational resources used to perform the data analysis were provided by ComputeCanada (www.computecanada.org) and CLUMEQ (www.clumeq.mcgill.ca), which is funded in part by NSERC (MRS), FQRNT, and McGill University.

